# Dorsal and ventral striatal functional connectivity shift with the medial frontal gyrus in internet gaming disorder: Potential mechanisms underlying addictive engagement

**DOI:** 10.1101/2020.08.16.253385

**Authors:** Guang-Heng Dong, Haohao Dong, Min Wang, Jialin Zhang, Weiran Zhou, Yanbin Zheng, Marc N. Potenza

**Affiliations:** Center for Cognition and Brain Disorders, the Affiliated Hospital of Hangzhou Normal University, Hangzhou, P.R. China; Institute of Psychological Research, Hangzhou Normal University, Hangzhou, P.R. China; Zhejiang Key Laboratory for Research in Assessment of Cognitive Impairments, Hangzhou, Zhejiang Province, P.R. China; Department of Psychology, Zhejiang Normal University, Jinhua, P.R. China; Department of Psychiatry and Child Study Center, Yale University School of Medicine, New Haven, CT, USA; Department of Neuroscience, Yale University, New Haven, CT, USA; Connecticut Council on Problem Gambling, Wethersfield, CT, USA; Connecticut Mental Health Center, New Haven, CT, USA

**Author notes:** Corresponding Authors: Guang-Heng Dong, Ph.D., Professor, Center for Cognition and Brain Disorders, The Affiliated Hospital of Hangzhou Normal University, 2318 Yuhangtang Road, Hangzhou, Zhejiang Province, PR China. E-Mail, Marc N. Potenza, Ph.D., M.D., Professor, Yale School of Medicine.

**Keywords:** Internet gaming disorder, striatum, functional connectivity, effective connectivity, longitudinal study

## Abstract

**Background:** Animal models suggest transitions from non-addictive to addictive behavioral engagement are associated with ventral-to-dorsal striatal shifts. However, few studies have examined such features in humans, especially in internet gaming disorder (IGD), a behavioral addiction.

**Methods:** Four-hundred-and-eighteen subjects (174 with IGD; 244 with recreational game use (RGU)) were recruited. Resting-state fMRI data were collected and functional connectivity (FC) analyses were performed based on ventral and dorsal striatal seeds. Correlations and follow-up spectrum dynamic causal model (spDCM) analyses were performed to examine relationships between ventral/dorsal striatum to medial frontal gyrus (MFG) and IGD severity. Longitudinal data from 40 subjects (22 IGD; 18 RGU) were also analysed to investigate further.

**Results:** Interactions were observed between group (IGD, RGU) and striatal regions (ventral, dorsal). IGD relative to RGU subjects showed lower ventral-striatum-to-MFG (mostly involving supplementary motor area (SMA)) and higher dorsal-striatum-to-MFG functional connectivity. spDCM revealed that left dorsal-striatum-to-MFG connectivity was correlated with IGD severity. Longitudinal data further support for ventral-to-dorsal striatal MFG relationships in IGD.

**Conclusions:** Consistent with animal models of substance addictions, ventral-to-dorsal striatal transitions in involvement coritico-striatal circuitry may underlie IGD and its severity. These findings suggest possible neurobiological mechanisms that may be targeted in treatments for IGD.

## Introduction

Altered frontal-striatal communication may relate to impaired control over motivations that promote engagement in and persistence of addictive behaviors (Fineberg et al., 2010; Zhou et al., 2018). The striatum, important in reward processing, has been implicated in transitions in addictions. The striatum also contributes to associative learning, motor function, behavioural control and other functions that may contribute to engagement in addictive behaviours (Contreras-Rodriguez, Martin-Perez, Vilar-Lopez, & Verdejo-Garcia, 2017). Changes in striatal functioning have thus been proposed as central elements in behavioral addictions like internet gaming disorder (IGD) (Brand, Young, Laier, Wolfling, & Potenza, 2016; Dong & Potenza, 2014; Zhou et al., 2018).

The striatum may be divided into ventral and dorsal components, based on functional and anatomical evidence, and different striatal subdivisions may have distinct roles (Contreras-Rodriguez et al., 2017; Zhou et al., 2018). The dorsal striatum (DS) may be particularly important for flexible versus habitual action selection and action, and the ventral striatum (VS) may be particularly important for learning values of stimuli (Balleine, Liljeholm, & Ostlund, 2009; Marche, Martel, & Apicella, 2017). Animal models have suggested that transitions from non-addictive to addictive engagement may involve shifts in involvement of ventral-to-dorsal striatal cortico-striato-thalamo-cortical circuits (Brand et al., 2019; Fineberg et al., 2010; Zhou et al., 2018).

Preclinical studies indicate that self-administration of addictive drugs, such as stimulants like cocaine, may lead to DS neuroadaptations (Everitt & Robbins, 2016; Smith & Robbins, 2013). Human neuroimaging studies of drug craving indicate that drug-related cues activate DS regions implicated in habitual behaviours and linked to measures of compulsivity in drug-using populations (Everitt & Robbins, 2016; Hyatt et al., 2012; Volkow et al., 2006). Therefore, DS neuroadaptations have been implicated in transitions between incentive-based and habit-based control of behaviours, and greater involvement of the DS is often observed when substance/drug intake is more addictive or compulsive (Contreras-Rodriguez et al., 2017; Volkow, Wang, Tomasi, & Baler, 2013).

Shifts in VS-DS function has recently been implicated in humans with cannabis (Zhou et al., 2018) or tobacco (Hu, Salmeron, Gu, Stein, & Yang, 2015) use disorders and those with obesity (Contreras-Rodriguez et al., 2017). Behavioural addictions such as gambling and gaming disorders do not involve substances linked to the development of maintenance of these disorders or their recoveries (Dong, Lin, Zhou, & Lu, 2014; Dong, Liu, Zheng, Du, & Potenza, 2019; G. H. Dong, M. Wang, H. Zheng, et al., 2020). Nonetheless, similar circuitry has been proposed to underlie such transitions (Brand et al., 2019). However, few studies have examined such proposed models directly.

Internet gaming disorder (IGD) has been included in DSM-5 as a possible condition warranting additional research (American_Psychiatric_Association, 2013). IGD is characterized by poorly controlled gaming that leads to impairment or psychological distress and persists despite negative consequences (G. H. Dong, M. Wang, Z. Wang, et al., 2020; Zhang et al., 2020). Classified as a behavioural addiction (Petry and O’Brien 2013; WHO 2017), IGD may involve impaired executive control (Z. Wang et al., 2019; Zheng et al., 2019), enhanced craving to gaming cues (Dong et al., 2019; Zhang et al., 2020), and disadvantageous decision-making (Dong & Potenza, 2014).

Although studies have investigated neurobiological correlates of IGD, a minority has used resting-state fMRI assessments that may permit evaluations of VS-to-DS shifts that are not influenced by tasks or experimental paradigms (A. Weinstein, Livny, & Weizman, 2017; A. M. Weinstein, 2017; Zheng et al., 2019). Resting-state functional connectivity (FC) facilitates assessment of functional alterations in the absence of contextual modulation (Di Martino et al., 2011; Di Martino et al., 2008). This approach has provided insight into frontostriatal functional circuits in cannabis (Blanco-Hinojo et al., 2017; Zimmermann et al., 2018) and tobacco (Hu et al., 2015) use disorders.

Many people play video games in a controlled manner and exhibit recreational game use (RGU) (Dong et al., 2019; G. Dong et al., 2020; G. H. Dong, M. Wang, Z. Wang, et al., 2020). In the current study, we used people with RGU as a control group, and by comparing IGD to RGU subjects, we sought to examine brain features associated with IGD and potential changes over time. Identification of such features could provide new insights into neurobiological mechanisms of IGD and suggest potential therapeutic targets for the disorder.

Altogether, while both VS and DS networks have been proposed to be relevant to stages of IGD and other addictions, including with respect to addiction severity, few studies have directly assessed these relationships, particularly over time. Here, we aimed to investigate VS and DS FC in IGD versus RGU subjects and examine individual differences therein with respect to IGD severity. To do so, we applied a seed-based approach to resting-state fMRI data to assess VS and DS FC. Resting-state fluctuations may relate to cognitive and emotional biases that may in part shape individual preferences; thus, striatal connectivity measures may have predictive validity in relation to IGD severity (Buckner & Vincent, 2007; Contreras-Rodriguez et al., 2017; C. G. Yan et al., 2019). We hypothesized that IGD versus RGU participants would show relatively increased FC in the DS, particularly with respect to cortical regions implicated IGD. We further hypothesized that increased DS FC would be associated with IGD severity, and these relationships would be observed longitudinally.

## Materials and methods

### Ethics

The experiment conforms to the Code of Ethics of the World Medical Association (Declaration of Helsinki). The Human Investigations Committee of Hangzhou Normal University approved this research. All subjects were university students from Shanghai and were recruited through advertisements. All participants provided written informed consent before experimentation.

### Participant selection

Valid resting-state data from 174 IGD subjects and 244 RGU subjects scanned in 2016-2019 were included in the current study. Exclusion criteria (e.g., incomplete information, poor spatial normalization or brain coverage, excessive head motion) and inclusion criteria are provided (Table 1).

Criteria for selection of IGD and RGU have been reported previously (G. Dong et al., 2020) and are described briefly below. IGD was determined based on scores of 50 or more on Young’s online internet addiction test (IAT, www.netaddiction.com) (Young, 2009) and concurrently meeting proposed DSM-5 IGD criteria (Petry et al., 2014) (Table 1). RGU participants were required to meet fewer than 5 (of 9) of the proposed DSM-5 criteria for IGD and score less than 50 on Young’s IAT.

All participants were right-handed and were university students recruited through advertisements. All participants provided written informed consent and underwent structured psychiatric interviews (using the Mini-International Neuropsychiatric Interview (MINI)) (Lecrubier et al., 1997) performed by an experienced psychiatrist. All participants were free of psychiatric disorders (including major depression, anxiety disorders, schizophrenia, and substance dependence disorders) as assessed by the MINI. Depression was further accessed with Beck Depression Inventory (BDI) and those who scored higher than 4 were excluded.

### Data acquisition

Resting-state functional data (T2*-weighted images) were acquired using a 3T Siemens Trio MRI scanner at the East China Normal University. Participants were instructed to keep their head still and eyes open during scanning. Earplugs and a head coil with foam pads were used to minimize machine noise and head motion. Specific parameters are as follows: repetition time = 2000ms, interleaved 33 axial slices, echo time = 30ms, thickness = 3.0 mm, flip angle = 90°, field of view (FOV) = 220mm × 220mm, matrix = 64 × 64. Participants kept their eyes closed and were instructed not think of anything in particular during scanning. In order to minimize head movement, all subjects’ heads were cushioned with foam padding. Each fMRI scan lasted 7min, consisting of 210 imaging volumes.

### Data pre-processing

Resting-state data analysis was performed using REST and DPARSF (http://restfmri.net) (C.-G. Yan & Zang, 2010). Preprocessing was performed using a standard approach that consisted of the following steps: (1) the initial 10 volumes were discarded, and slice-timing correction was performed; (2) the time series of images for each subject were realigned using a six-parameter (rigid body) linear transformation; (3) individual T1-weighted images were co-registered to the mean functional image using a 6 degrees-of-freedom linear transformation without re-sampling and then segmented into gray matter (GM), white matter (WM) and cerebrospinal fluid (CSF); (4) transformations from individual native space to MNI space were computed with the DARTEL tool; (5) head-motion scrubbing using the Friston 24-parameter model to regress out head-motion effects; (6) mean framewise displacement (derived from Jenkinson’s relative root mean square algorithm) was used to address the residual effects of motion as a covariate in group analyses; (7) further preprocessing included band-pass filtering between 0.01 and 0.08 Hz.

### Region of interest (ROI) selection and seed-to-voxel functional connectivity

Anatomical seeds were chosen to increase the generalizability of findings (Duncan, 2013; Mansouri, Tanaka, & Buckley, 2009). We selected the bilateral NAcc (ventral striatum), the bilateral putamen and caudate (dorsal striatum) based on the HarvardOxford-sub-maxprob-thr25 atlas.

For each seed, FC was acquired by Pearson correlation coefficients between the mean time series of the seed region and all brain voxels (defined by the binary GM mask in SPM). A Fisher’s Z transformation was applied to improve normality of correlation coefficient values. Finally, two-sample *t*-tests were applied to map group differences of connectivity maps for each seed between IGD and RGU participants.

### Group-level image processing

For group-level functional connectivity analyses, we performed two-sample *t*-tests to compare the connectivity maps of each seed between IGD and RGU participants. Corrections for multiple comparisons were conducted using permutation-based inferences (5,000 permutations) with Threshold-Free Clustering Enhancement (TFCE), which provides strict control while improving replicability (Chen, Lu, & Yan, 2018). TFCE is a strict multiple-comparison correction strategy, which has been described as reaching the best balance between family-wise error rate (under 5%) and test-retest reliability/replicability relative to AFNI 3dClustSim, DPABI AlphaSim GRF, and false discovery rate (Chen et al., 2018). For regions showing significant group differences (*t-*test), the mean Fisher Z-transformed correlation coefficients were extracted from the underlying anatomical regions for further post-hoc analysis.

### FC results with gaming history, symptom severity

We also correlated changes in functional connectivity with gaming history and addiction severity among IGD individuals. A *P*<0.05 threshold was considered as significant.

### Spectral dynamic causal modelling (spDCM)

To provide insight into possible causal relatiionships between correlated brain regions, we performed spDCM analyses (implemented by DCM12.5 (revision 7487), which is based on SPM12 (https://www.fil.ion.ucl.ac.uk/spm/)). A detailed description of the DCM specification can be found in Friston et al (2003; 2014) and our previous work (G. H. Dong et al., 2020; M. Wang, Zheng, Du, & Dong, 2019). Briefly, we first constructed a general linear model for each participant’s data, and extracted and corrected the principal eigenvariate from the selected ROIs. A single “full” model was then specified for each participant. Here, the “full” model means that a two-way influence is assumed between regions. Then, we completed the estimation of the first-level DCM model by inverting the “full” models.

### Replication test with longitudinal data

To investigate further, we tracked 40 subjects (22 IGD, 18 RGU) for more than 6 months and obtained additional data. These 40 subjects was part of the 418 subjects. We collected their resting state data and addiction features during these two times of scans. The details of these 40 subjects were listed in Supplementary Table 2. All pre-processing steps, group-level image processing and functional connectivity calculations are of the same with the cross-sectional analyses.

## Results

### Ventral and dorsal features with brain regions

When examining the dorsal and ventral striatal ROIs, significant interactions were observed in relation to MFG connectivity in IGD and RGU subjects. Further analyses showed that IGD subjects showed relatively greater MFG-DS versus MFG-VS FC than RGU subjects in both hemispheres. (Figure 1).

**Figure 1:**
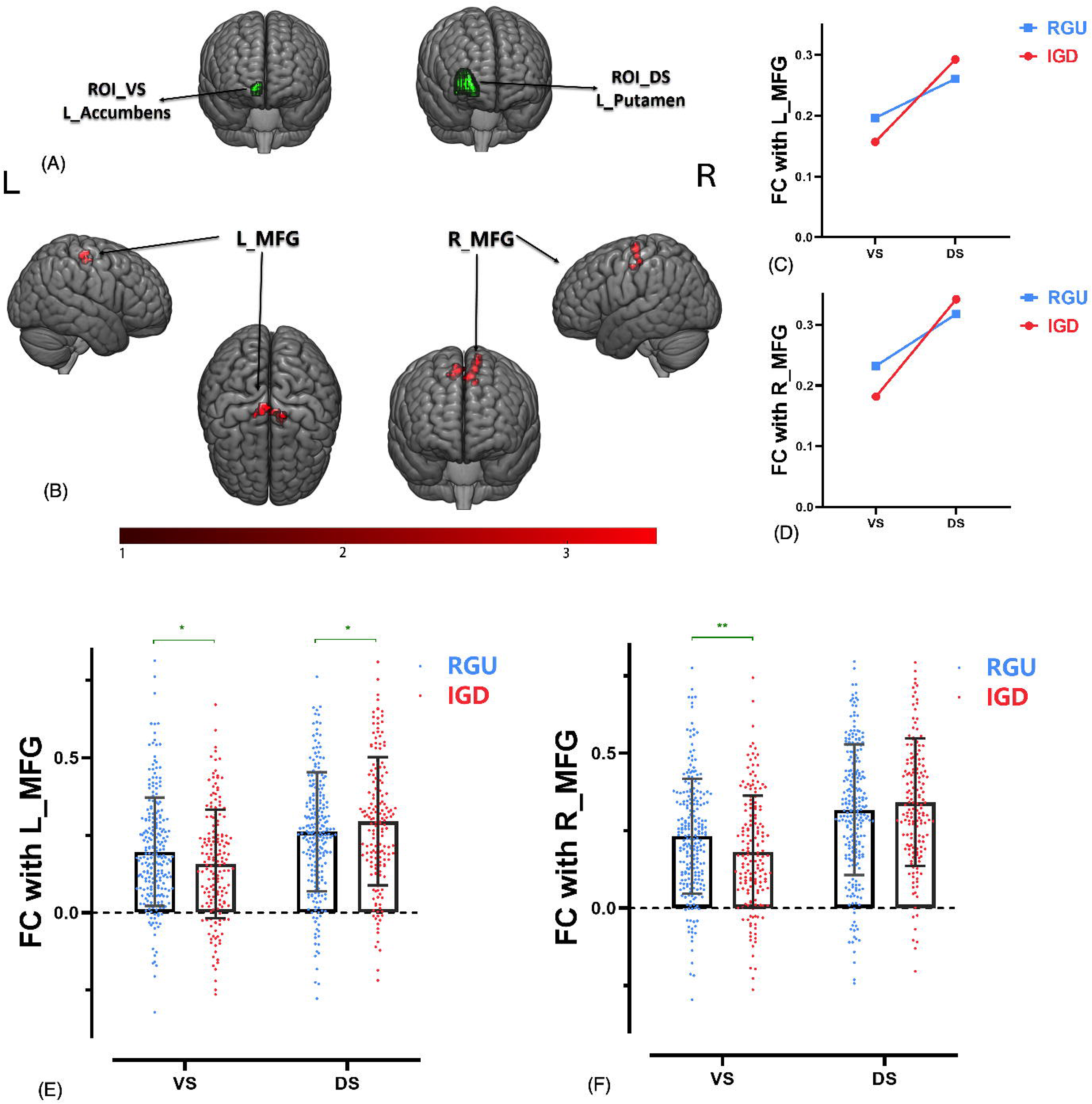
Interactions were found between ventral/dorsal striatum and the MFG. A: ROIs selected for the ventral and dorsal striatum in the current study. Left hemispheric regions are shown. B: Brain regions that show FC interactions with the ventral/dorsal striatum seeds and diagnostic groups. C, D: Plots demonstrating the interaction on FC strength between ventral/dorsal striatum and left and right MFG in IGD and RGU subjects. E,F: Strength of the FC between the VS/DS and left or right MFG in IGD and RGU subjects

### Correlation with behavioral results

A significant correlation was observed between left DS-MFG communication and the DSM-5 scores in IGD subjects. No correlation was observed in RGU subjects (Figure 2). No correlations were observed in the VS-MFG FC with DSM-5 scores.

**Figure 2:**
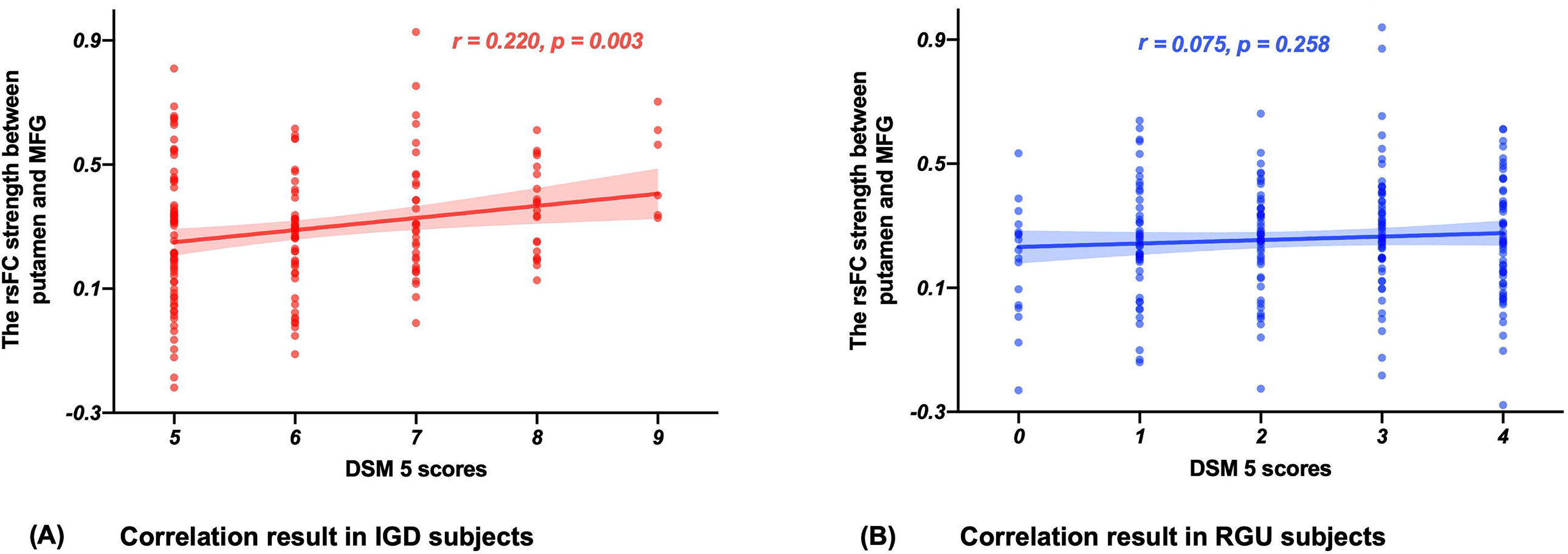
Correlations between IGD severity (DSM-5 score) and FC strength between left putamen (DS) and MFG. A: The correlation results in IGD subjects. B: The correlation results in RGU subjects.

### spDCM results

Dynamic causal modelling results showed a significant left MFG-to-DS (Putamen) communication. Specifically, with increasing addiction severity, the MFG-to-DS FC was stronger (Figure 3).

**Figure 3:**
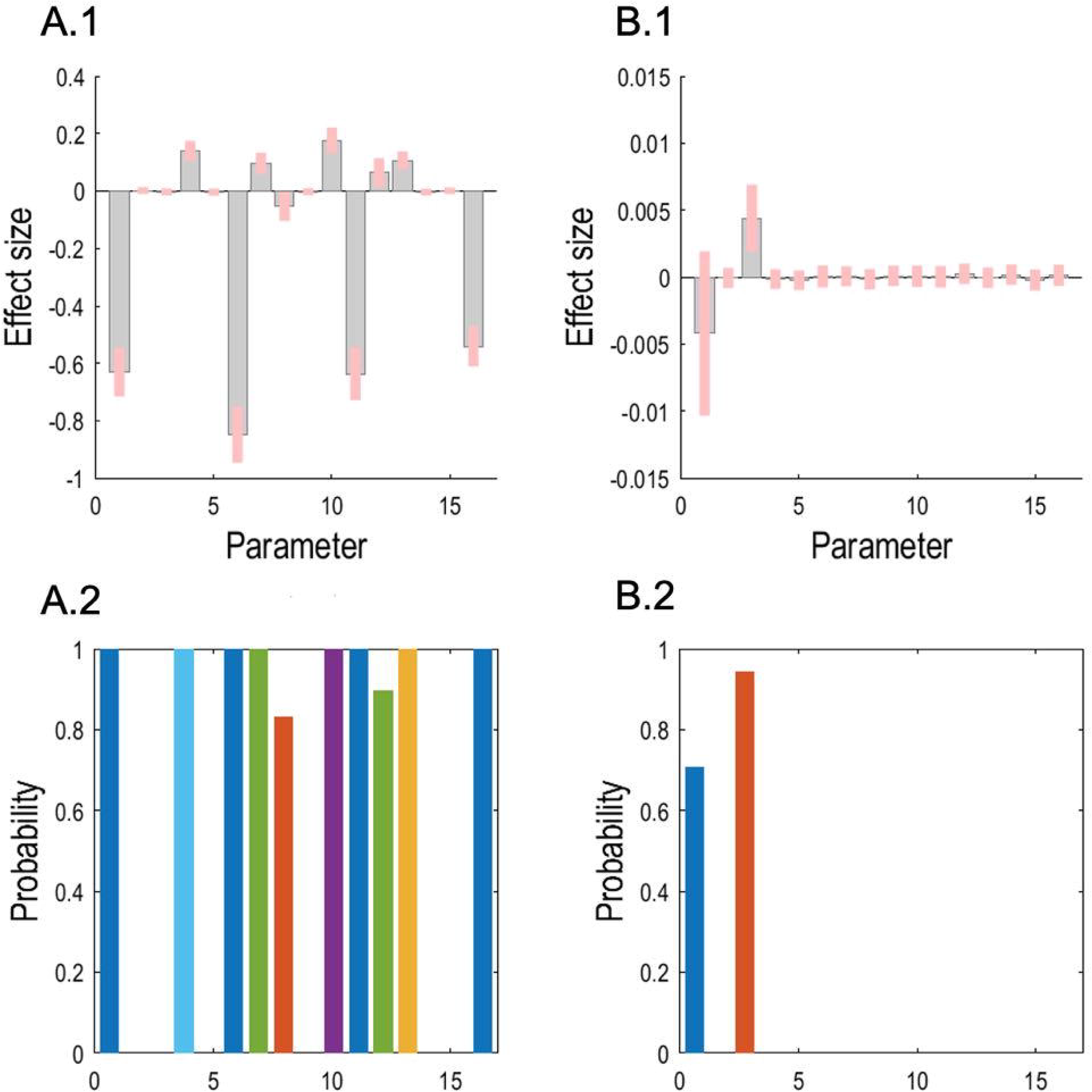
Correlation results between effective connections and IAT scores through parametric empirical Bayesian framework. A.1 shows each parameter’s effect size after Bayesian model averaging for the mean of all subjects, and A.2 shows the corresponding posterior probabilities. B.1 shows the effect size of correlations between each parameter and IAT scores and B.2 shows the corresponding posterior probabilities. Here the threshold of the posterior probability is set to 0.9, so the current results indicate that a positive correlation between connection strength and IAT is observed in effective connection from the MFG to left putamen (i.e., parameter 3 in B.1, B.2).

### Longitudinal data

Similar VS-to-DS features were observed in first and second scan among a small sample followed longitudinally (Figure 4). Over time in IGD subjects, VS-to-DS features in IGD may further strengthen. This relationship was not observed in RGU subjects.

**Figure 4:**
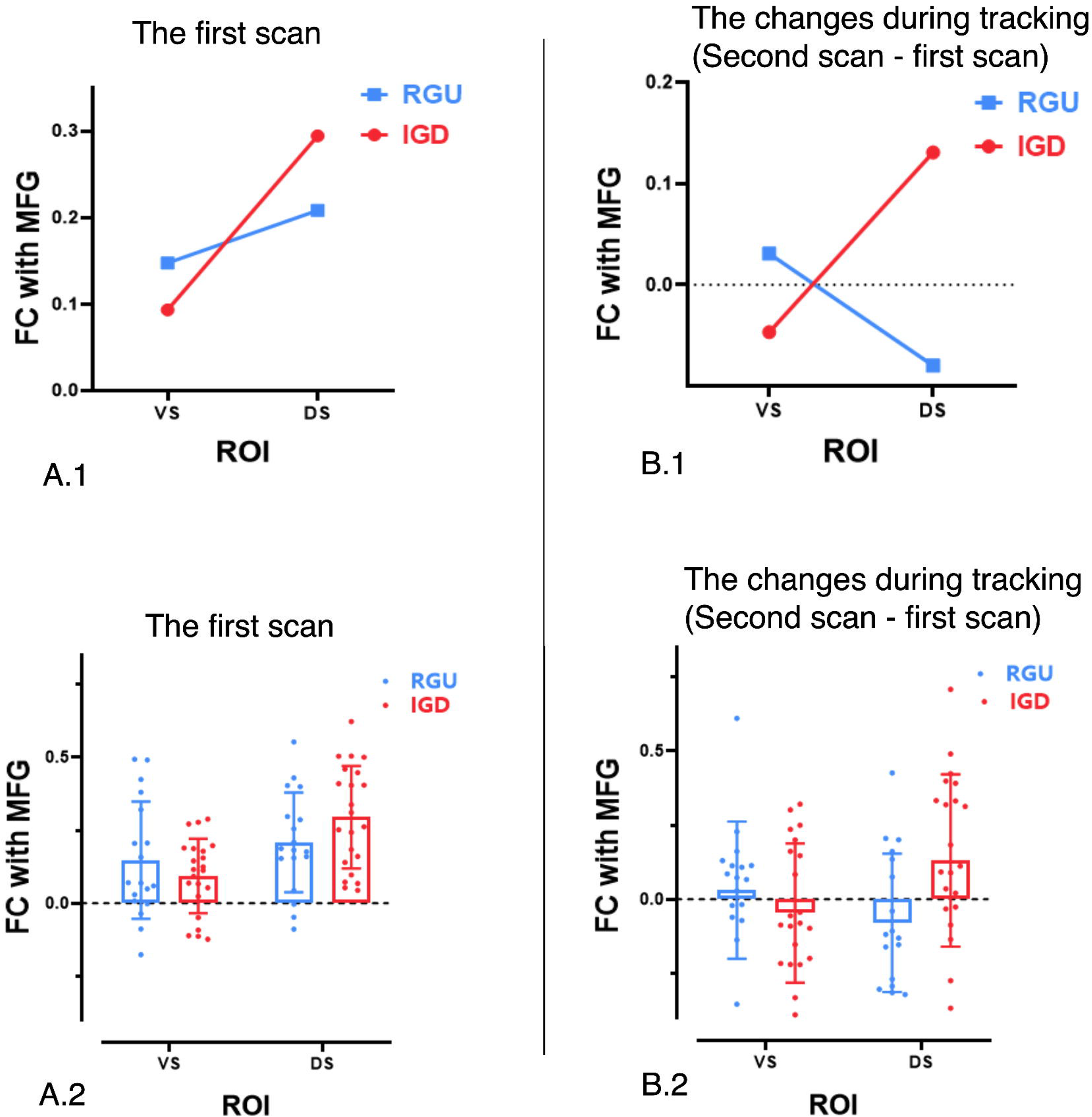
The VS-to-DS involvement in longitudinal data. A.1, A.2: The FC strength between ventral/dorsal striatum and MFG in IGD and RGU groups in the first scan. B.1, B.2: The FC strength changes between ventral/dorsal striatum and MFG in IGD and RGU groups in the following scan relative to the first scan.

### Rigor/reproducibility

We assessed rigor and reproducibility by testing whether global signal regression (GSR) influenced findings. With GSR, we observed similar results. Although the size of clusters varied slightly, similar patterns were found (See supplementary Figure 1-3; Supplementary table 1). This confirmation indicating that the results appear stable and not affected by GSR is important as studies have suggested that global signal may reflect spurious noise (Power, Plitt, Laumann, & Martin, 2017).

## Discussion

Using selected ROIs, the current study examined FC in the DS and VS in IGD and RGU subjects. We first identified an interactive effect between diagnostic group (IDG, RGU) and seed region (VS, DS) in FC with the MFG a large sample. Next, we demonstrated similar results longitudinally in a sub-group of the subjects. Results are consonant in suggesting the relevance of DS versus VS connectivity to the MFG in IGD, raising the possibility of VS to DS shifts in IGD that have been proposed. The current results deepen our understanding of neural features underlying IGD and suggests possible brain targets for neuromodulation interventions.

### Ventral striatum and MFG

In the VS, IGD relative to RGU subjects showed lower FC with bilateral MFG (mostly located in the supplementary motor area (SMA)). The SMA has been implicated in motor planning and execution, including self-initiated movements, action monitoring, response inhibition and action sequencing (Bonini et al., 2014; Cona & Semenza, 2017; Mostofsky & Simmonds, 2008; Passingham, Bengtsson, & Lau, 2010). The SMA may “link cognition to action” (Nachev, Kennard, & Husain, 2008) and act as a hub for processes related to motor intentionality (Albertini et al., 2020; Zapparoli et al., 2018). The motor basal-ganglia-thalamocortical circuit model proposes that the circuit originates at the SMA, receiving input from primary and the premotor cortices (Alexander, DeLong, & Strick, 1986). These areas connect with the nuclei of the basal ganglia, which project back to the thalamus, closing the loop by reconnecting with the site of origin in motor cortical areas (Alexander et al., 1986; Owens-Walton et al., 2019).

As the VS contributes to learning values of stimuli (Balleine et al., 2009), relatively decreased FC between the SMA and VS in IGD suggests a possible disconnection between stimulus evaluation and behavioural/motoric responses in domains including response inhibition. While currently speculative, this notion could be examined further by investigating the degree to which such FC relates to aspects of poor impulse control or other potentially related features in IGD. The current study complements previous ones indicating neural correlates of: gaming cues that typically elicit higher cravings in IGD versus RGU subjects (G. Dong et al., 2020; G. H. Dong, M. Wang, H. Zheng, et al., 2020); reward and loss processing that suggest that IGD subjects may show enhanced sensitivity to reward and decreased sensitivity to punishment (Dong, Huang, & Du, 2011; Dong et al., 2018); and, inter-temporal decision-making in which IGD subjects preferentially select smaller, sooner over larger, later rewards (Y. Wang, Hu, et al., 2017; Y. Wang, Wu, et al., 2017; Zheng et al., 2019). Together, results suggest possible complex neural mechanisms underlying features of IGD.

### Dorsal striatum and MFG

IGD relative to RGU subjects show greater DS-to-SMA FC. FC between the DS and SMA was observed in multiple studies. Individuals with first-episode psychosis compared to those without have shown greater FC between the left dorsal caudate and left motor cortex (Oh, Kim, Kim, Lee, & Kwon, 2019). FC between the DS and sensorimotor cortex is important for action–reward associations (Balleine, Delgado, & Hikosaka, 2007; Balleine et al., 2009), and this has been suggested as a pathophysiological mechanism of motivational deficits in schizophrenia (Stepien et al., 2018). Increased connectivity between the DS and motor cortex has also been linked to social anhedonia (Y. Wang et al., 2016). These findings suggest that relatively increased FC between the striatum and motor cortex may relate to negative mood states, and these have often been linked to IGD and related behaviours (Desai, Krishnan-Sarin, Cavallo, & Potenza, 2010; Liu, Desai, Krishnan-Sarin, Cavallo, & Potenza, 2011). As the current study largely excluded individuals with features of depression, additional research is needed to investigate whether DS-MFG FC may be stronger in individuals with IGD and co-occurring depression. However, significant correlations between IGD severity and DS-MFG FC suggest this relationship may underlie other clinically relevant aspects of IGD. The findings are consistent with theorical models of IGD (Brand et al., 2019; Brand et al., 2016; Dong & Potenza, 2014) that proposes that IGD is associated with neural adaptations in the striatum, particularly with DS networks as IGD becomes more established and engrained.

Drug addictions have also been linked to neuroadaptations in the DS (Balleine et al., 2009; Marche et al., 2017). Thus, across behavioural and drug addictions, DS-cortical circuitry may importantly relate to habitual behaviours. The extent to which DS-MFG FC relates to measures of habit and compulsivity in IGD warrants further direct examination. The DCM results suggest that the greater the DS-to-MFG communication, the more severe the IGD. The longitudinal sample extends the findings to indicate that this findings persists over time. Future studies should examine the extent to which possible shifts in VS-to-DS FC with the MFG relate to progression of more severe IGD, and greater DS-to-VS FC with the MFG may relate to recovery. The extent to which this circuitry may be targeted by neuromodulation warrants direct examination, especially given the accessibility of the MFG to non-invasive transcranial manipulation.

### Limitations

Several limitations should be noted. First, more men than women were recruited in current study, which limits exploration of possible gender-related differences. Second, the study involved young adults from China and the extent to which the findings extend to other jurisdictions and cultures warrants additional investigation.

### Conclusions

In summary, the findings that IGD is linked to relatively greater DS-MFG FC suggests a link between more habitual and poorly controlled gaming in IGD to a ventral-to-dorsal shift in cortico-striatal circuitry involvement. Future studies investigating the malleability of the connectivity of such circuits to interventions are needed.

## Acknowledgements

The current research was supported by the Zhejiang Provincial Natural Science Foundation (LY20C090005). Marc Potenza received support from the Connecticut Council on Problem Gambling, the Connecticut Department of Mental Health and Addictions and the National Center for Responsible Gaming. The funding agencies did not have input into the writing of this manuscript.

## Authors’ contributions

Guang-Heng Dong designed this research and wrote the first draft of the manuscript. Haohao Dong analysed the data and prepared the figures and tables on pre-processing and the interactions between VS and DS, and the longitudinal data analyses; Min Wang analysed the spDCM results and prepared relevant figures. Jialin Zhang, Weiran Zhou, Yanbin Zheng contributed to fMRI data collection. Marc Potenza edited the manuscript. All authors contributed to and approved the final manuscript.

## References

Albertini, D., Gerbella, M., Lanzilotto, M., Livi, A., Maranesi, M., Ferroni, C. G., & Bonini, L. (2020).Connectional gradients underlie functional transitions in monkey pre-supplementary motor area. Prog Neurobiol, 184, 101699. doi:10.1016/j.pneurobio.2019.101699s

Alexander, G. E., DeLong, M. R., & Strick, P. L. (1986).Parallel organization of functionally segregated circuits linking basal ganglia and cortex. Annu Rev Neurosci, 9, 357–381. doi:10.1146/annurev.ne.09.030186.002041

American_Psychiatric_Association. (2013). Diagnostic and statistical manual of mental disorders (5th ed.).

Balleine, B. W., Delgado, M. R., & Hikosaka, O. (2007).The role of the dorsal striatum in reward and decision-making. J Neurosci, 27(31), 8161–8165. doi:10.1523/JNEUROSCI.1554-07.2007

Balleine, B. W., Liljeholm, M., & Ostlund, S. B. (2009).The integrative function of the basal ganglia in instrumental conditioning. Behav Brain Res, 199(1), 43–52. doi:10.1016/j.bbr.2008.10.034

Blanco-Hinojo, L., Pujol, J., Harrison, B. J., Macia, D., Batalla, A., Nogue, S., … Martin-Santos, R. (2017).Attenuated frontal and sensory inputs to the basal ganglia in cannabis users. Addict Biol, 22(4), 1036–1047. doi:10.1111/adb.12370

Bonini, F., Burle, B., Liegeois-Chauvel, C., Regis, J., Chauvel, P., & Vidal, F. (2014).Action monitoring and medial frontal cortex: leading role of supplementary motor area. Science, 343(6173), 888–891. doi:10.1126/science.1247412

Brand, M., Wegmann, E., Stark, R., Muller, A., Wolfling, K., Robbins, T. W., & Potenza, M. N. (2019).The Interaction of Person-Affect-Cognition-Execution (I-PACE) model for addictive behaviors: Update, generalization to addictive behaviors beyond internet-use disorders, and specification of the process character of addictive behaviors. Neurosci Biobehav Rev, 104, 1–10. doi:10.1016/j.neubiorev.2019.06.032

Brand, M., Young, K. S., Laier, C., Wolfling, K., & Potenza, M. N. (2016).Integrating psychological and neurobiological considerations regarding the development and maintenance of specific Internet-use disorders: An Interaction of Person-Affect-Cognition-Execution (I-PACE) model. Neuroscience and Biobehavioral Reviews, 71, 252–266. doi:10.1016/j.neubiorev.2016.08.033

Buckner, R. L., & Vincent, J. L. (2007).Unrest at rest: default activity and spontaneous network correlations. Neuroimage, 37(4), 1091-1096; discussion 1097-1099. doi:10.1016/j.neuroimage.2007.01.010

Chen, X., Lu, B., & Yan, C. G. (2018).Reproducibility of R-fMRI metrics on the impact of different strategies for multiple comparison correction and sample sizes. Hum Brain Mapp, 39(1), 300–318. doi:10.1002/hbm.23843

Cona, G., & Semenza, C. (2017). Supplementary motor area as key structure for domain-general sequence processing: A unified account. Neurosci Biobehav Rev, 72, 28–42. doi:10.1016/j.neubiorev.2016.10.033

Contreras-Rodriguez, O., Martin-Perez, C., Vilar-Lopez, R., & Verdejo-Garcia, A. (2017). Ventral and Dorsal Striatum Networks in Obesity: Link to Food Craving and Weight Gain. Biol Psychiatry, 81(9), 789–796. doi:10.1016/j.biopsych.2015.11.020

Desai, R. A., Krishnan-Sarin, S., Cavallo, D., & Potenza, M. N. (2010). Video-gaming among high school students: health correlates, gender differences, and problematic gaming. Pediatrics, 126(6), e1414–1424. doi:10.1542/peds.2009-2706

Di Martino, A., Kelly, C., Grzadzinski, R., Zuo, X. N., Mennes, M., Mairena, M. A., … Milham, M. P. (2011). Aberrant striatal functional connectivity in children with autism. Biol Psychiatry, 69(9), 847–856. doi:10.1016/j.biopsych.2010.10.029

Di Martino, A., Scheres, A., Margulies, D. S., Kelly, A. M., Uddin, L. Q., Shehzad, Z., … Milham, M. P. (2008). Functional connectivity of human striatum: a resting state FMRI study. Cereb Cortex, 18(12), 2735–2747. doi:10.1093/cercor/bhn041

Dong, G., Huang, J., & Du, X. (2011). Enhanced reward sensitivity and decreased loss sensitivity in Internet addicts: an fMRI study during a guessing task. J Psychiatr Res, 45(11), 1525–1529. doi:10.1016/j.jpsychires.2011.06.017

Dong, G., Lin, X., Zhou, H., & Lu, Q. (2014). Cognitive flexibility in internet addicts: fMRI evidence from difficult-to-easy and easy-to-difficult switching situations. Addict Behav, 39(3), 677–683. doi:10.1016/j.addbeh.2013.11.028

Dong, G., Liu, X., Zheng, H., Du, X., & Potenza, M. N. (2019). Brain response features during forced break could predict subsequent recovery in internet gaming disorder: A longitudinal study. J Psychiatr Res, 113, 17–26. doi:10.1016/j.jpsychires.2019.03.003

Dong, G., & Potenza, M. N. (2014). A cognitive-behavioral model of Internet gaming disorder: theoretical underpinnings and clinical implications. J Psychiatr Res, 58, 7–11. doi:10.1016/j.jpsychires.2014.07.005

Dong, G., Wang, M., Liu, X., Liang, Q., Du, X., & Potenza, M. N. (2020). Cue-elicited craving-related lentiform activation during gaming deprivation is associated with the emergence of Internet gaming disorder. Addict Biol, 25(1), e12713. doi:10.1111/adb.12713

Dong, G., Zheng, H., Liu, X., Wang, Y., Du, X., & Potenza, M. N. (2018). Gender-related differences in cue-elicited cravings in Internet gaming disorder: The effects of deprivation. Journal of Behavioral Addictions, 1-12. doi:10.1556/2006.7.2018.118

Dong, G. H., Wang, M., Wang, Z., Zheng, H., Du, X., & Potenza, M. N. (2020). Addiction severity modulates the precuneus involvement in internet gaming disorder: Functionality, morphology and effective connectivity. Prog Neuropsychopharmacol Biol Psychiatry, 98, 109829. doi:10.1016/j.pnpbp.2019.109829

Dong, G. H., Wang, M., Zheng, H., Wang, Z., Du, X., & Potenza, M. N. (2020). Disrupted prefrontal regulation of striatum-related craving in Internet gaming disorder revealed by dynamic causal modeling: results from a cue-reactivity task. Psychol Med, 1–13. doi:10.1017/S003329172000032X

Duncan, J. (2013). The Structure of Cognition: Attentional Episodes in Mind and Brain. Neuron, 80(1), 35–50. doi:10.1016/j.neuron.2013.09.015

Everitt, B. J., & Robbins, T. W. (2016). Drug Addiction: Updating Actions to Habits to Compulsions Ten Years On. Annu Rev Psychol, 67, 23–50. doi:10.1146/annurev-psych-122414-033457

Fineberg, N. A., Potenza, M. N., Chamberlain, S. R., Berlin, H. A., Menzies, L., Bechara, A., … Hollander, E. (2010). Probing Compulsive and Impulsive Behaviors, from Animal Models to Endophenotypes: A Narrative Review. Neuropsychopharmacology, 35(3), 591–604. doi:10.1038/npp.2009.185

Hu, Y., Salmeron, B. J., Gu, H., Stein, E. A., & Yang, Y. (2015). Impaired functional connectivity within and between frontostriatal circuits and its association with compulsive drug use and trait impulsivity in cocaine addiction. JAMA Psychiatry, 72(6), 584–592. doi:10.1001/jamapsychiatry.2015.1

Hyatt, C. J., Assaf, M., Muska, C. E., Rosen, R. I., Thomas, A. D., Johnson, M. R., … Pearlson, G. D. (2012). Reward-related dorsal striatal activity differences between former and current cocaine dependent individuals during an interactive competitive game. Plos One, 7(5), e34917. doi:10.1371/journal.pone.0034917

Lecrubier, Y., Sheehan, D. V., Weiller, E., Amorim, P., Bonora, I., Harnett Sheehan, K., … Dunbar, G. C. (1997). The Mini International Neuropsychiatric Interview (MINI). A short diagnostic structured interview: reliability and validity according to the CIDI. European Psychiatry, 12(5), 224–231.

Liu, T. C., Desai, R. A., Krishnan-Sarin, S., Cavallo, D. A., & Potenza, M. N. (2011). Problematic Internet use and health in adolescents: data from a high school survey in Connecticut. J Clin Psychiatry, 72(6), 836–845. doi:10.4088/JCP.10m06057

Mansouri, F. A., Tanaka, K., & Buckley, M. J. (2009). Conflict-induced behavioural adjustment: a clue to the executive functions of the prefrontal cortex. Nature Reviews Neuroscience, 10(2), 141–152. doi:10.1038/nrn2538

Marche, K., Martel, A. C., & Apicella, P. (2017). Differences between Dorsal and Ventral Striatum in the Sensitivity of Tonically Active Neurons to Rewarding Events. Front Syst Neurosci, 11, 52. doi:10.3389/fnsys.2017.00052

Mostofsky, S. H., & Simmonds, D. J. (2008). Response inhibition and response selection: two sides of the same coin. J Cogn Neurosci, 20(5), 751–761. doi:10.1162/jocn.2008.20500

Nachev, P., Kennard, C., & Husain, M. (2008). Functional role of the supplementary and pre-supplementary motor areas. Nat Rev Neurosci, 9(11), 856–869. doi:10.1038/nrn2478

Oh, S., Kim, M., Kim, T., Lee, T. Y., & Kwon, J. S. (2019). Resting-state functional connectivity of the striatum predicts improvement in negative symptoms and general functioning in patients with first-episode psychosis: A 1-year naturalistic follow-up study. Aust N Z J Psychiatry, 4867419885452. doi:10.1177/0004867419885452

Owens-Walton, C., Jakabek, D., Power, B. D., Walterfang, M., Velakoulis, D., van Westen, D., … Hansson, O. (2019). Increased functional connectivity of thalamic subdivisions in patients with Parkinson’s disease. Plos One, 14(9), e0222002. doi:10.1371/journal.pone.0222002

Passingham, R. E., Bengtsson, S. L., & Lau, H. C. (2010). Medial frontal cortex: from self-generated action to reflection on one’s own performance. Trends Cogn Sci, 14(1), 16–21. doi:10.1016/j.tics.2009.11.001

Petry, N. M., Rehbein, F., Gentile, D. A., Lemmens, J. S., Rumpf, H. J., Mossle, T., … O’Brien, C. P. (2014). An international consensus for assessing internet gaming disorder using the new DSM-5 approach. Addiction, 109(9), 1399–1406. doi:10.1111/add.12457

Power, J. D., Plitt, M., Laumann, T. O., & Martin, A. (2017). Sources and implications of whole-brain fMRI signals in humans. Neuroimage, 146, 609–625. doi:10.1016/j.neuroimage.2016.09.038

Smith, D. G., & Robbins, T. W. (2013). The neurobiological underpinnings of obesity and binge eating: a rationale for adopting the food addiction model. Biol Psychiatry, 73(9), 804–810. doi:10.1016/j.biopsych.2012.08.026

Stepien, M., Manoliu, A., Kubli, R., Schneider, K., Tobler, P. N., Seifritz, E., … Kirschner, M. (2018). Investigating the association of ventral and dorsal striatal dysfunction during reward anticipation with negative symptoms in patients with schizophrenia and healthy individuals. Plos One, 13(6), e0198215. doi:10.1371/journal.pone.0198215

Volkow, N. D., Wang, G. J., Telang, F., Fowler, J. S., Logan, J., Childress, A. R., … Wong, C. (2006). Cocaine cues and dopamine in dorsal striatum: mechanism of craving in cocaine addiction. J Neurosci, 26(24), 6583–6588. doi:10.1523/JNEUROSCI.1544-06.2006

Volkow, N. D., Wang, G. J., Tomasi, D., & Baler, R. D. (2013). The addictive dimensionality of obesity. Biol Psychiatry, 73(9), 811–818. doi:10.1016/j.biopsych.2012.12.020

Wang, Y., Hu, Y., Xu, J., Zhou, H., Lin, X., Du, X., & Dong, G. (2017). Dysfunctional Prefrontal Function Is Associated with Impulsivity in People with Internet Gaming Disorder during a Delay Discounting Task. Front Psychiatry, 8, 287. doi:10.3389/fpsyt.2017.00287

Wang, Y., Liu, W. H., Li, Z., Wei, X. H., Jiang, X. Q., Geng, F. L., … Chan, R. C. (2016). Altered corticostriatal functional connectivity in individuals with high social anhedonia. Psychol Med, 46(1), 125–135. doi:10.1017/S0033291715001592

Wang, Y., Wu, L., Wang, L., Zhang, Y., Du, X., & Dong, G. (2017). Impaired decision-making and impulse control in Internet gaming addicts: evidence from the comparison with recreational Internet game users. Addict Biol, 22(6), 1610–1621. doi:10.1111/adb.12458

Wang, Z., Liu, X., Hu, Y., Zheng, H., Du, X., & Dong, G. (2019). Altered brain functional networks in Internet gaming disorder: independent component and graph theoretical analysis under a probability discounting task. CNS Spectr, 24(5), 544–556. doi:10.1017/S1092852918001505

Weinstein, A., Livny, A., & Weizman, A. (2017). New developments in brain research of internet and gaming disorder. Neurosci Biobehav Rev, 75, 314–330. doi:10.1016/j.neubiorev.2017.01.040

Weinstein, A. M. (2017). An Update Overview on Brain Imaging Studies of Internet Gaming Disorder. Front Psychiatry, 8, 185. doi:10.3389/fpsyt.2017.00185

Yan, C.-G., & Zang, Y.-F. (2010). DPARSF: A MATLAB Toolbox for “Pipeline” Data Analysis of Resting-State fMRI. Front Syst Neurosci, 4, 13. doi:10.3389/fnsys.2010.00013

Yan, C. G., Chen, X., Li, L., Castellanos, F. X., Bai, T. J., Bo, Q. J., … Zang, Y. F. (2019). Reduced default mode network functional connectivity in patients with recurrent major depressive disorder. Proc Natl Acad Sci U S A, 116(18), 9078–9083. doi:10.1073/pnas.1900390116

Young, K. S. (2009, Jan. 2010). Internet Addiction Test (IAT). Retrieved from http://netaddiction.com/index.php?option=combfquiz&view=onepage&catid=46&Itemid=106

Zapparoli, L., Seghezzi, S., Scifo, P., Zerbi, A., Banfi, G., Tettamanti, M., & Paulesu, E. (2018). Dissecting the neurofunctional bases of intentional action. Proc Natl Acad Sci U S A, 115(28), 7440–7445. doi:10.1073/pnas.1718891115

Zhang, J., Dong, H., Zhao, Z., Chen, S., Jiang, Q., Du, X., & Dong, G. H. (2020). Altered neural processing of negative stimuli in people with internet gaming disorder: fMRI evidence from the comparison with recreational game users. J Affect Disord, 264, 324–332. doi:10.1016/j.jad.2020.01.008

Zheng, H., Hu, Y., Wang, Z., Wang, M., Du, X., & Dong, G. (2019). Meta-analyses of the functional neural alterations in subjects with Internet gaming disorder: Similarities and differences across different paradigms. Prog Neuropsychopharmacol Biol Psychiatry, 94, 109656. doi:10.1016/j.pnpbp.2019.109656

Zhou, F., Zimmermann, K., Xin, F., Scheele, D., Dau, W., Banger, M., … Becker, B. (2018). Shifted balance of dorsal versus ventral striatal communication with frontal reward and regulatory regions in cannabis-dependent males. Hum Brain Mapp, 39(12), 5062–5073. doi:10.1002/hbm.24345

Zimmermann, K., Yao, S., Heinz, M., Zhou, F., Dau, W., Banger, M., … Becker, B. (2018). Altered orbitofrontal activity and dorsal striatal connectivity during emotion processing in dependent marijuana users after 28 days of abstinence. Psychopharmacology (Berl), 235(3), 849–859. doi:10.1007/s00213-017-4803-6

