## Supplementary material for "Dorsal and ventral striatal functional connectivity shift with the medial frontal gyrus in internet gaming disorder: Potential mechanisms underlying addictive engagement": Table 1

Table 1. Demographics and clinical characteristics of all subjects

|  | IGD | RGU | Group difference |
| --- | --- | --- | --- |
|  | N=174  Male=102  Female =72 | N=244  Male=149  Female=95 | *p*-value |
| Demographics (M±SD) | |  |  |
| Age (year) | 21.20±2.441 | 21.56±2.478 | .156 |
| Years of education | 14.57±1.474 | 14.61±1.402 | .823 |
| Clinical characteristics (M±SD) | |  |  |
| IAT | 65.72±8.783 | 40.09±10.571 | <.001 |
| DSM | 6.12±1.145 | 2.53±1.430 | <.001 |
| Gaming time (hours/week) | 8.27±3.616 | 6.17±3.024 | <.001 |
| Gaming history (year) | 3.72±.640 | 3.73±.698 | .894 |
| craving | 51.79±17.304 | 34.77±16.280 | <.001 |

**Abbreviations:** IGD, Internet Gaming Disorder; RGU, recreational game use; M, mean; SD, standard deviation; IAT, Internet Addiction Test; DSM, number of DSM-5 item
