## Supplementary material for "Dorsal and ventral striatal functional connectivity shift with the medial frontal gyrus in internet gaming disorder: Potential mechanisms underlying addictive engagement": Table 2

**Table 2: Results that show functional connectivity interactions between the VS and DS in the current study**

| ROI | cluster | Peak MNI coordinates | | | Peak Intensity | Cluster Size | Region | AAL |
| --- | --- | --- | --- | --- | --- | --- | --- | --- |
|  |  | X | Y | Z |  |  |  |  |
| Left  Putamen- Accumbens | 1 | 12 | -15 | 63 | 4.58 | 55 | R Medial Frontal Gyrus | Supp_Motor_Area_R |
|  | 2 | -9 | -24 | 57 | 4.3246 | 37 | L Medial Frontal Gyrus | Supp_Motor_Area_L |

All images are thresholded at *p* < .05, TFCE-corrected; Iterations =5000.

MNI: Montreal Neurological Institute; AAL: Anatomical Automatic Labelling; TFCE:Threshold-Free Clustering Enhancement.
