## supplementary materials for "Dorsal and ventral striatal functional connectivity shift with the medial frontal gyrus in internet gaming disorder: Potential mechanisms underlying addictive engagement"

**Rigor/Reproducibility**

We assessed rigor and reproducibility by testing whether global signal regression (GSR) influenced the results. With GSR, we found similar results. Although the size of the clusters varied slightly, similar conclusions can be drawn (See Supplementary Figure 1-3; Supplementary Table 1).

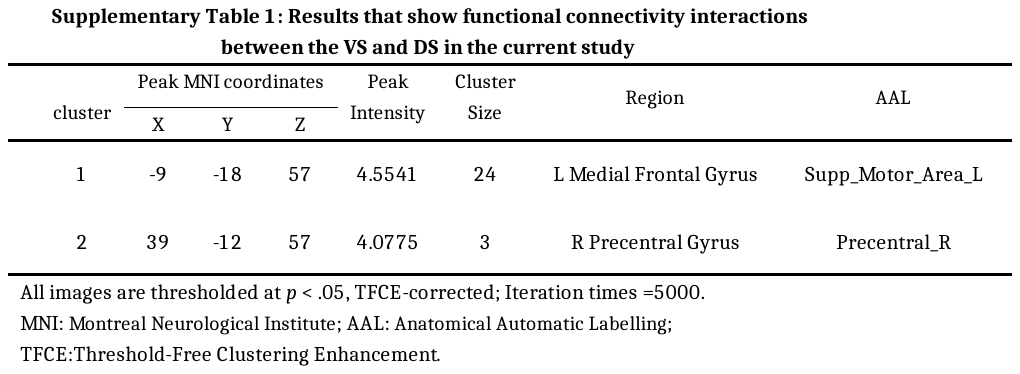

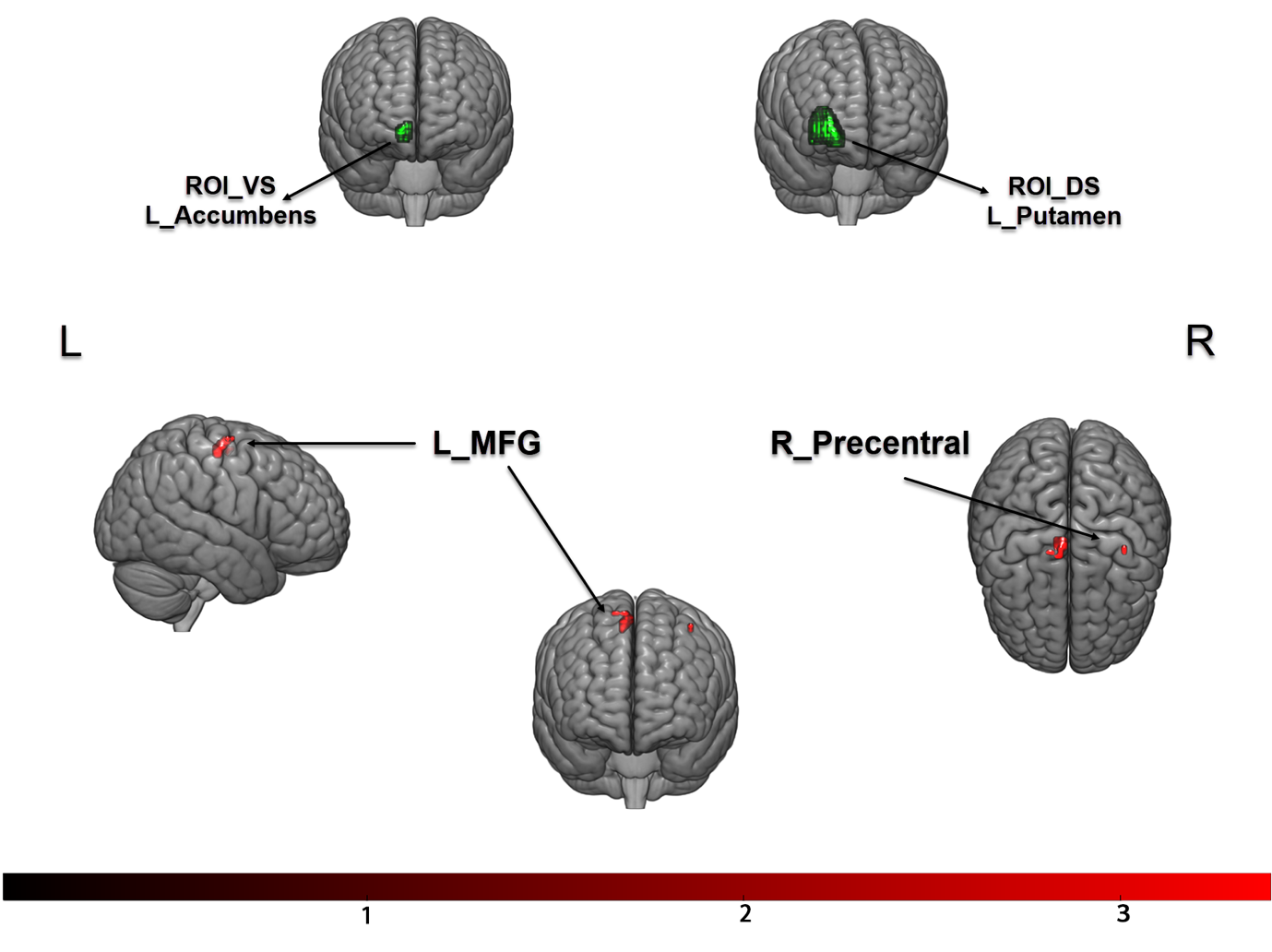

**Supplementary Figure 1: Interactions were found between ventral/dorsal striatum and MFG**

Upper: ROIs selected for ventral and dorsal in current study

Bottom: Brain regions that show FC interactions with left ventral/dorsal striatum

No significant results were observed in the right striatum.

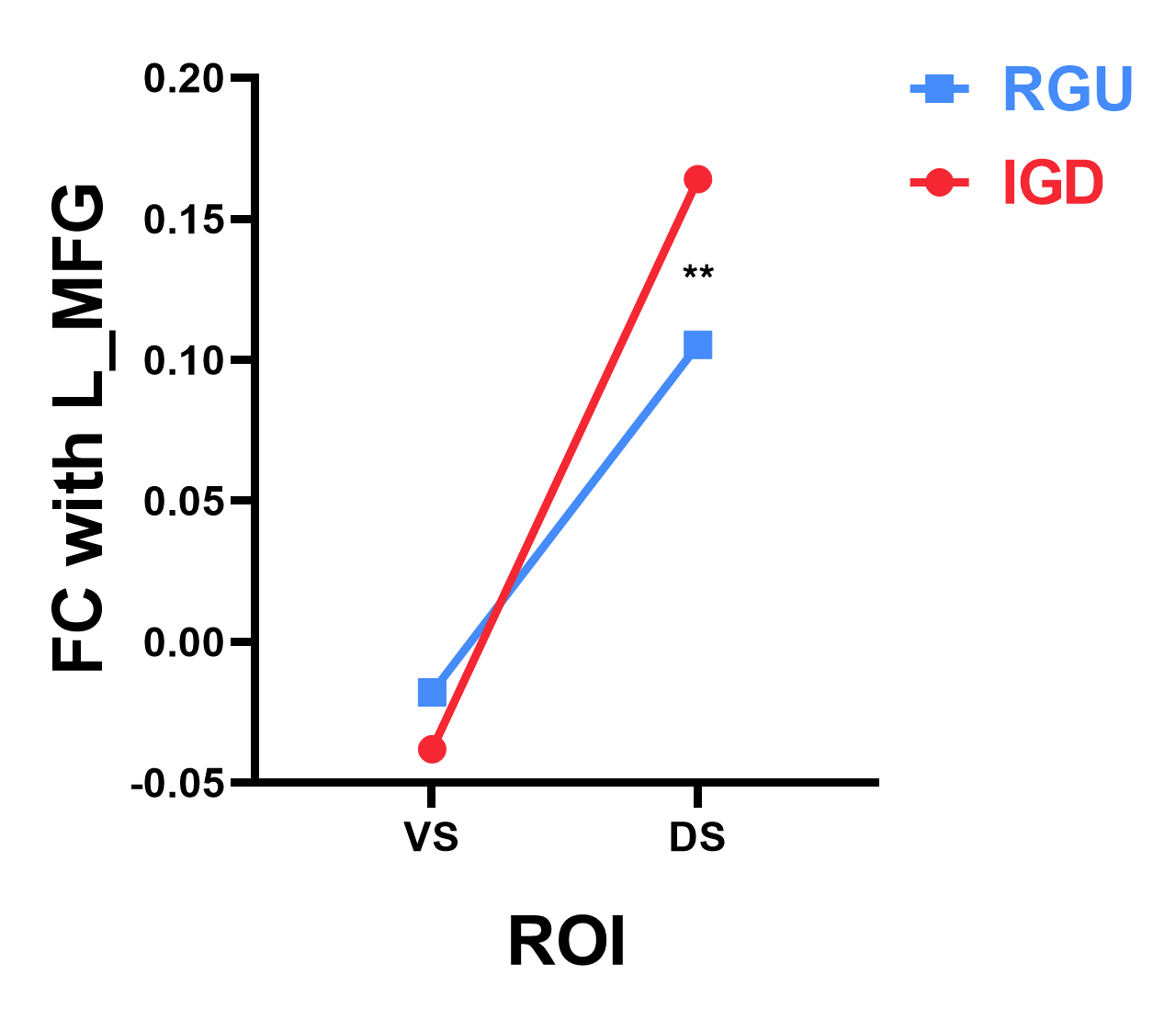

**Supplementary Figure 2:** **The FC between ventral/dorsal striatum and MFG in different groups.**

** *p*<0.01

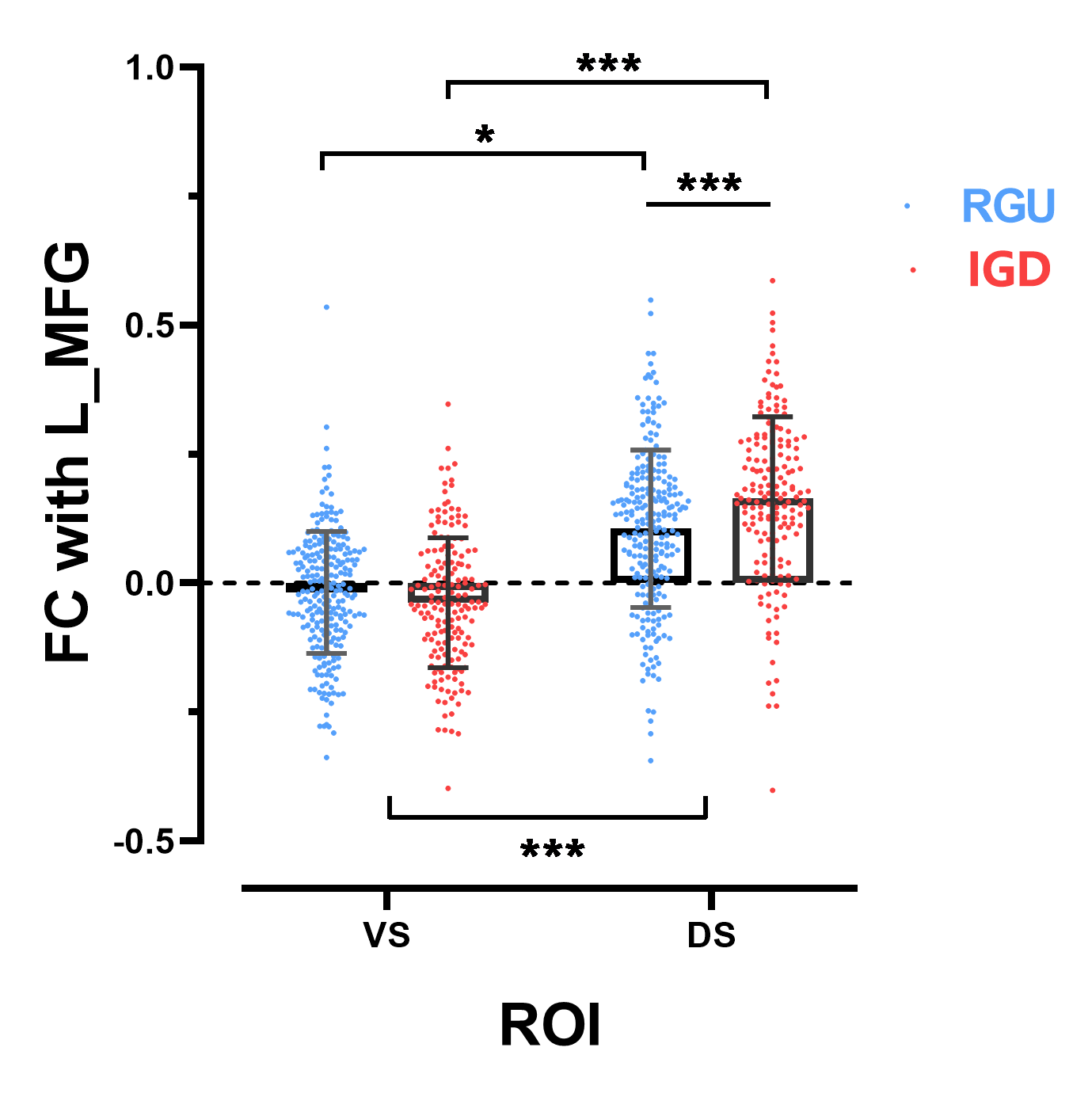

**Supplementary Figure 3: Detailed features between VS/DS and MFG in different comparisons**

The features in VS/DS FC with left MFG. * *p*<0.05; *** *p*<0.001

**Supplementary Table 2. Demographic information of 40 tracked subjects**

|  | | **IGD (n=22)**  **12 males** | **RGU (n=18)**  **9 males** | ***t*** | ***p*** |
| --- | --- | --- | --- | --- | --- |
| Age (years) | pre- | *20.4±2.0* | *20.2±1.9* | *0.277* | *0.783* |
|  | post- | *21.1±2.1* | *21.1±1.8* | *0.039* | *0.969* |
| Interval (months) |  | *9.2±3.5* | *9.3±3.6* | *-0.053* | *0.958* |
| IAT | pre- | *67.6±8.9* | *40.5±7.0* | *11.732* | *<0.001* |
|  | post- | *68.7±7.9* | *36.7±8.0* | *11.402* | *<0.001* |
| DSM-5 | pre- | *6.2±1.1* | *2.4±1.1* | *10.218* | *<0.001* |
|  | post- | *6.5±1.3* | *2.3±1.4* | *9.180* | *<0.001* |
| Game time (hours per week) | pre- | *26.9±9.2* | *13.7±4.5* | *5.849* | *<0.001* |
|  | post- | *27.6±10.3* | *17.0±8.4* | *3.500* | *0.001* |
| Craving | pre- | *53.0±19.3* | *31.7±16.3* | *3.711* | *0.001* |
|  | post- | *54.9±14.7* | *33.8±14.5* | *4.528* | *<0.001* |

IGD: Internet gaming disorder; RGU: recreational game user; IAT: Internet addiction test; DSM-5: Diagnostic and Statistical Manual of Mental Disorders-5; BDI: Beck Depression Inventory.
